# The *Helicobacter pylori* TlpD cytoplasmic chemoreceptor requires an intact C-terminus for polar localization and function

**DOI:** 10.1101/2024.11.08.622596

**Authors:** Raymondo Lopez-Magaña, Karen M. Ottemann

## Abstract

Bacteria localize proteins to distinct subcellular locations, including chemoreceptors, which frequently localize to the bacterial pole. Although some polarity-promoting mechanisms have been described, many chemoreceptors lack clear routes to becoming polar. TlpD of the bacterial pathogen *Helicobacter pylori* is one such protein. This cytoplasmic chemoreceptor localizes to the pole in a manner that is independent of the other chemoreceptors. In this work, we evaluated the role of TlpD domains in its function. Truncated proteins were created that lacked different amounts of the N- or C-termini and expressed in *H. pylori* in place of native *tlpD* or as the sole chemoreceptor. These TlpD variants were examined for expression, protein localization, association with chemotaxis signaling proteins, and effect on motility. TlpD that lacked any portion of the N-terminal 104 amino acids did not produce detectable protein. In contrast, TlpD retained expression with loss of the C-terminal 45 amino acids. TlpD lacking the last 45 amino acids (TlpDΔC4) preserved the ability to interact with CheW and CheV proteins based on bacterial two-hybrid analysis but were unable to localize to the pole either on their own or in the presence of other chemoreceptors. TlpDΔC4 was found diffuse in the cytoplasm, and interacted with CheV1, CheV2, and CheV3 at this location but not CheW. TlpDΔC4 did not confer chemotactic abilities in soft agar chemotaxis assays. These findings suggest the C-terminal end of TlpD plays a previously unappreciated role in promoting TlpD polar localization and function.

**Importance:** Bacteria place their proteins in specific locations that are required for the proteins to function including the bacterial pole. How the bacterial cell identifies which proteins go to the pole is not fully understood. In this work, we dissect parts of a protein called TlpD that naturally goes to the pole. We find that mutants lacking one end of TlpD lose their polar placement, but retain other abilities. TlpD allows directed motility known as chemotaxis. This ability is critical for infection in *H. pylori* and numerous other pathogens. When TlpD loses its polar placement, the protein no longer functions for chemotaxis, laying the foundation for future studies that can dissect how this segment promotes function and eventually translate into therapies for *H. pylori* infection.

## Introduction

*H. pylori* is a human pathogen that infects approximately one-half of the world’s population (1). This bacterium infects the stomach and causes chronic stomach inflammation and gastric cancer (1). To reach the stomach, *H. pylori* must navigate a dynamic environment. En route, *H. pylori* encounters various signals, including mammalian produced ones such as acid from stomach parietal cells and reactive oxygen from epithelial and innate immune cells (2–5). One strategy *H. pylori* uses to circumvent such harmful compounds and find beneficial ones is directed motility, also known as chemotaxis (6). The chemotaxis signal transduction system, extensively studied in *Escherichia coli*, allows bacteria to sense concentration gradients of particular chemicals and initiate a motility response (7–10). This chemotaxis system is required for *H. pylori* to establish infection (2, 11–14). The ability to sense and mount an appropriate response towards these signals allows effective navigation and infection.

The *H. pylori* chemotaxis system, like the *E. coli* one, operates as a multiprotein complex. Like that of well-studied bacteria, this complex forms via protein-protein interactions, including between sensors known as chemoreceptors that form interactions with coupling proteins and the CheA histidine kinase, to control kinase activity (15–17). In contrast to the one-coupling protein system of *E. coli* that uses the canonical coupling protein CheW, the chemotaxis system of *H. pylori* contains three additional coupling proteins called CheV1, CheV2, and CheV3 (15, 18). These CheV paralogs are proposed to function in several ways, including as coupling proteins, phosphate sinks, or possibly as unique coupling proteins to accommodate chemoreceptors that cannot couple to CheW (15, 19, 20). In the *H. pylori* system, both CheW and CheV1 are critical to build chemoreceptor arrays (16, 17) and each CheV plays specific roles in motility (18, 21). However, the exact roles of CheV2 and CheV3 in *H. pylori* chemotaxis have yet to be determined (15, 16).

The *H. pylori* chemotaxis system uses four chemoreceptors called TlpA, TlpB, TlpC, and TlpD. TlpA, TlpB and TlpC are integral membrane proteins whereas TlpD is soluble, not membrane bound and cytoplasmic (6, 22). This cytoplasmic chemoreceptor contains a C-terminal chemoreceptor zinc binding (CZB) sensing domain (17, 23). Despite being cytoplasmic, TlpD localizes to the pole and retains this capability even in the absence of any membrane bound chemoreceptors (17). However, the mechanism by which TlpD locates to the cell pole remains unclear.

TlpD recruits other chemotaxis proteins to the cell pole. This chemoreceptor is sufficient to recruit CheA, CheW, CheV1, and CheV3 proteins to the *H. pylori* cell pole (17). TlpD likely interacts with the CheW and CheV coupling proteins via its methyl accepting (MA) domain, as has been shown in other chemoreceptors. TlpD interacts directly with coupling proteins CheW and CheV1 (16). This interaction with coupling proteins facilitates an indirect interaction with the CheA kinase to form a functional signaling unit. Indeed, a combination of reconstituted TlpD, CheA and CheW or CheV1 proteins are sufficient to drive CheA phosphorylation and suggest either coupling protein can work with TlpD to create a functional signaling unit in vitro (16, 24). Moreover, *H. pylori* with TlpD as its only chemoreceptor retains spreading motility on soft agar, indicative of chemotaxis function, and thus supports a model by which TlpD alone forms an autonomous and functional signaling unit (17).

TlpD plays a key role in *H. pylori* infection. Numerous studies have shown that TlpD is required for full stomach colonization in both mice and gerbils (2, 11, 25). Of the four *H. pylori* chemoreceptors, *tlpD* mutants colonize less well than wild type or any isogenic chemoreceptor mutant (2, 11, 25). Additionally, TlpD mounts a chemotactic response to reactive oxygen species including hydrogen peroxide, superoxide and hypochlorous acid (17, 24). As all these signals are found during infection, it makes sense that TlpD would be an important chemoreceptor. Indeed, TlpD was found to be expressed in all tested *H. pylori* strains, suggesting it is retained to a higher degree than other chemoreceptors (17). These findings highlight the importance of TlpD in the chemotaxis system of *H. pylori* for infection.

In this study, we evaluated the roles of parts of TlpD for which there is no known function. We created TlpD truncations lacking portions of either the N or C terminus and analyzed how loss of these sequences affected function in situ, expressing the proteins in the native *H. pylori*. Most truncations resulted in loss of stability in *H. pylori* but one mutant construct, however, was stably expressed in *H. pylori*: a C-terminus deletion of the last amino acids, 388-433. This region is C-terminal to the CZB domain and based on bioinformatic analysis, had no detectable features or domains. This truncated protein, called TlpDΔC4, maintained its ability to interact with CheW and CheV coupling proteins, based on immunolocalization and bacterial two hybrid assays, but lost its ability to localize to the pole and function.

## Results

### Identification of TlpD domain endpoints to allow selective omission of unannotated regions

Previous work had identified domains of TlpD (23, 24) but to define precise endpoints, we carried out up-to-date bioinformatic analyses. AlphaFold2 (26) predicted the known MA and CZB domains (23) and provided a predicted structure of the 433 amino acid TlpD protein (Fig. 1A). Alphafold2 defined the MA domain as amino acids 152 to 287 and the CZB domain from 315 to 379 (Fig. 1B). Following the C-terminal CZB domain was a 54 amino acid stretch with no annotated domains (Fig. 1A and B). The N-terminus comprised the first 140 amino acids and is predicted to orient perpendicular to the rest of the protein (Fig. 1A and B). Like the C-terminus, the N-terminal stretch had no annotated motifs (Fig. 1A). Further analysis using the bioinformatic servers MotifFinder, NCBI CDD, Protein CCD similarly did not identify motifs in the N- or C-terminal regions (27–29) (Fig. S1 and File S1). Secondary structure analysis using Globplot (30) determined that these regions, along with most of TlpD, were structured (Fig. S1). Overall, these analyses suggest that TlpD has known domains in the middle portion, while the N- and C-termini are predicted to be folded into secondary structures but any lack identifiable domains.

**Figure 1.**
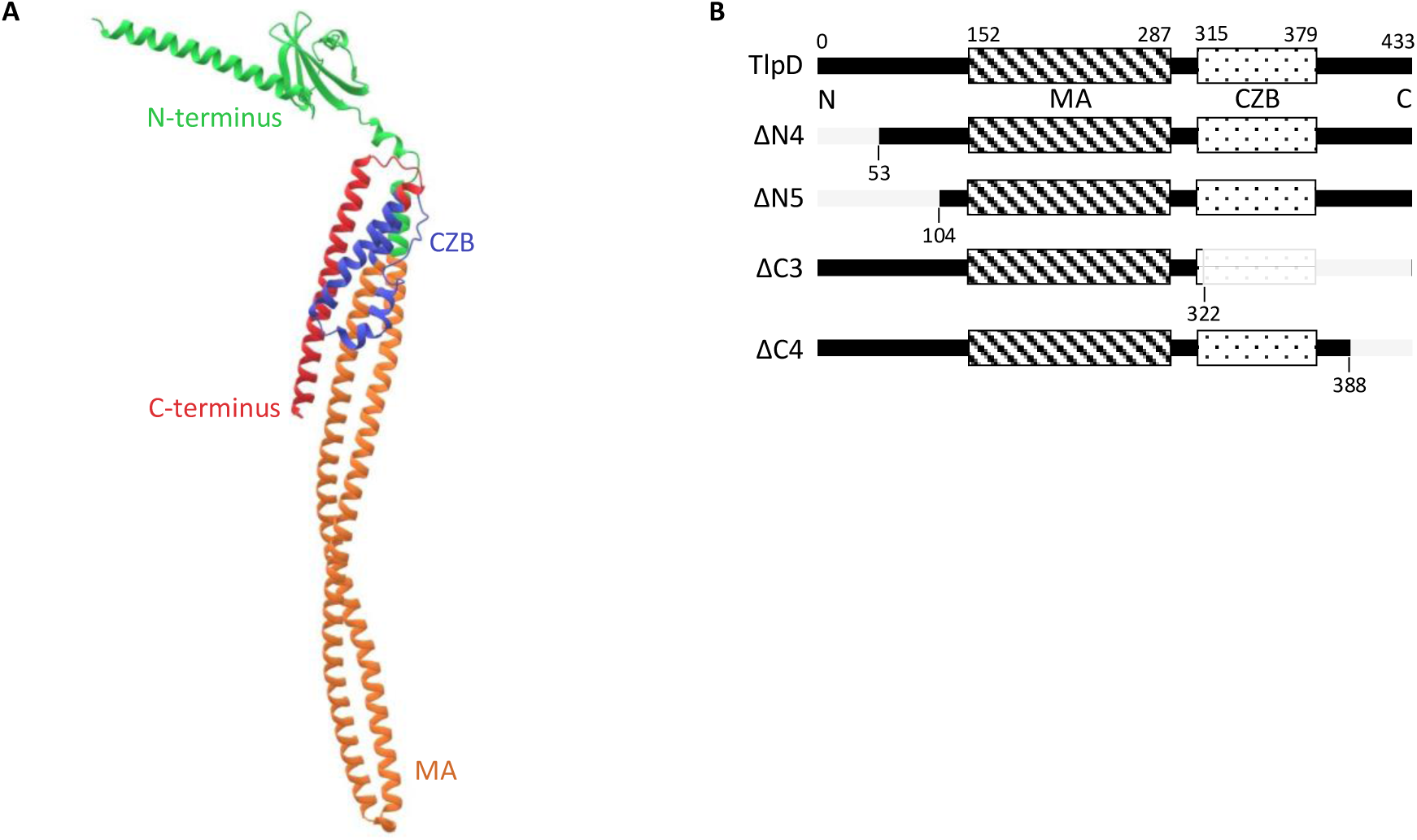
*H. pylori* TlpD predicted structure and domains. (A) AlphaFold predicted structure of *H. pylori* TlpD. TlpD is 433 amino acids in length with two known domains, the MA domain (orange) and CZB domain (blue). The region with no known domains at the N-terminus is shown in green, and at the C-terminus in red. (B) Diagram of modified versions of *H. pylori* TlpD with domains and truncation sites indicated (grey). TlpD represents the full-length version of the protein at 433 amino acids, ΔN4 is truncated at residue 53, ΔN5 is truncated at residue 104, ΔC3 is truncated at residue 322, and ΔC4 is truncated at residue 388. Each of these constructs contains a C-terminal 3xFLAG tag.

Because it is unclear what the functions of the TlpD N- or C-terminus are, we took advantage of the two-domain architecture of TlpD to create truncated versions that removed select regions but retained the MA or CZB domains (Fig. 1B). An initial round of constructs did not carefully avoid predicted secondary structure endpoints and resulted in proteins that were not expressed in *H. pylori* (Fig. S2). In the second round, predicted secondary structure endpoints were avoided and a 3X FLAG tag was added to improve detection (File S1 and S2). We designed these constructs to integrate into the non-essential *rdxA* locus of *H. pylori*, facilitated by an erythromycin resistance marker for positive selection and flanking *rdxA* sequences (Fig. S3A), in single copy under the control of the *tlpD* promoter (Table 1).

**Table 1.**
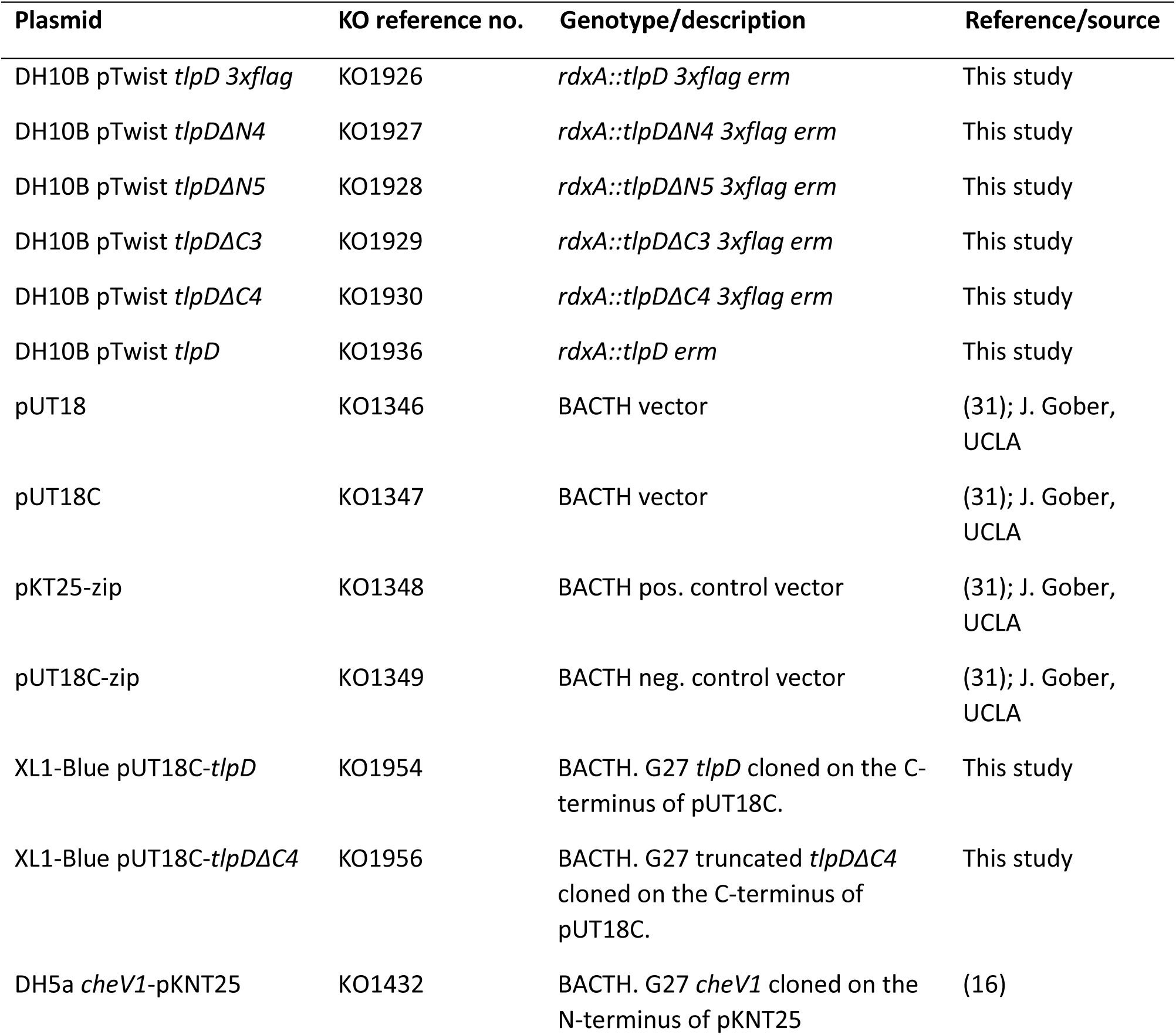
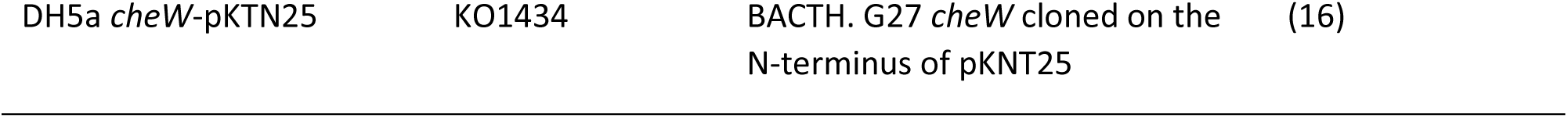
Plasmids used in this study.

### TlpD tolerates C-terminal but not the N-terminal truncations

To evaluate the production of these TlpD truncation constructs in vivo, each *tlpD* construct was transformed into *H. pylori* G27 *ΔtlpD* (Table 2), where they integrated into the genome at the *rdxA* locus (Fig. S3A). Proper integration was confirmed by PCR using primers that flank the *rdxA* locus (Fig. S3B). We then evaluated protein expression in *H. pylori* whole cell lysates by western blotting using an anti-FLAG antibody, as used in *E. coli* (Fig. 2). For this approach, *H. pylori* strains were grown on standard solid media (CHBA), scraped from the plate, normalized to a standard OD_600_ and lysed in Laemmli buffer. Full-length TlpD (TlpD) was detected when expressed from the *rdxA* locus (Fig. 2), but strains expressing the truncated variants showed variable expression. N-terminus truncated proteins (TlpD ΔN4 and ΔN5) had little or no protein detected (Fig. 2). Strains expressing C-terminus truncated proteins (TlpD ΔC3 and ΔC4) varied in the amounts of protein detected: TlpDΔC3 had a low signal, similar to TlpDΔN5, while TlpDΔC4 had a clearly detectable signal that was approximately half that of WT TlpD (Fig. 2). These findings suggest the C-terminal end of TlpD can be removed with modest effect on protein expression, and this protein was chosen for further characterization.

**Figure 2.**
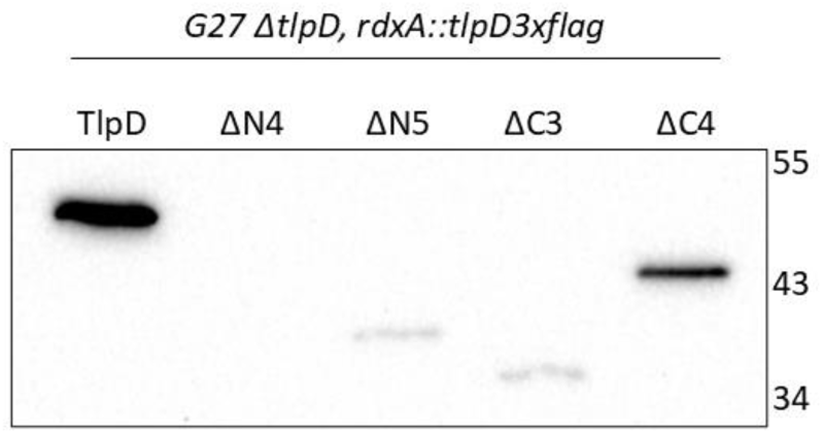
Analysis of expression of TlpD WT and truncated variants expressed from *H. pylori rdxA*. Western blot of *H. pylori* G27 Δ*tlpD* strains complemented with *tlpD* constructs were analyzed from whole-cell lysates with an anti-FLAG antibody, which recognizes the C-terminal tag. The TlpD variant expressed is indicated at the top, with TlpD indicating full length with the 3XFLAG. Marker sizes are given in kilodaltons on the right. The expected size of TlpD is 51 kDa, ΔN4 is 45 kDa, ΔN5 is 39 kDa, ΔC3 is 38 kDa, and ΔC4 is 45 kDa. Western blot is representative of 3 biological replicates.

**Table 2.**
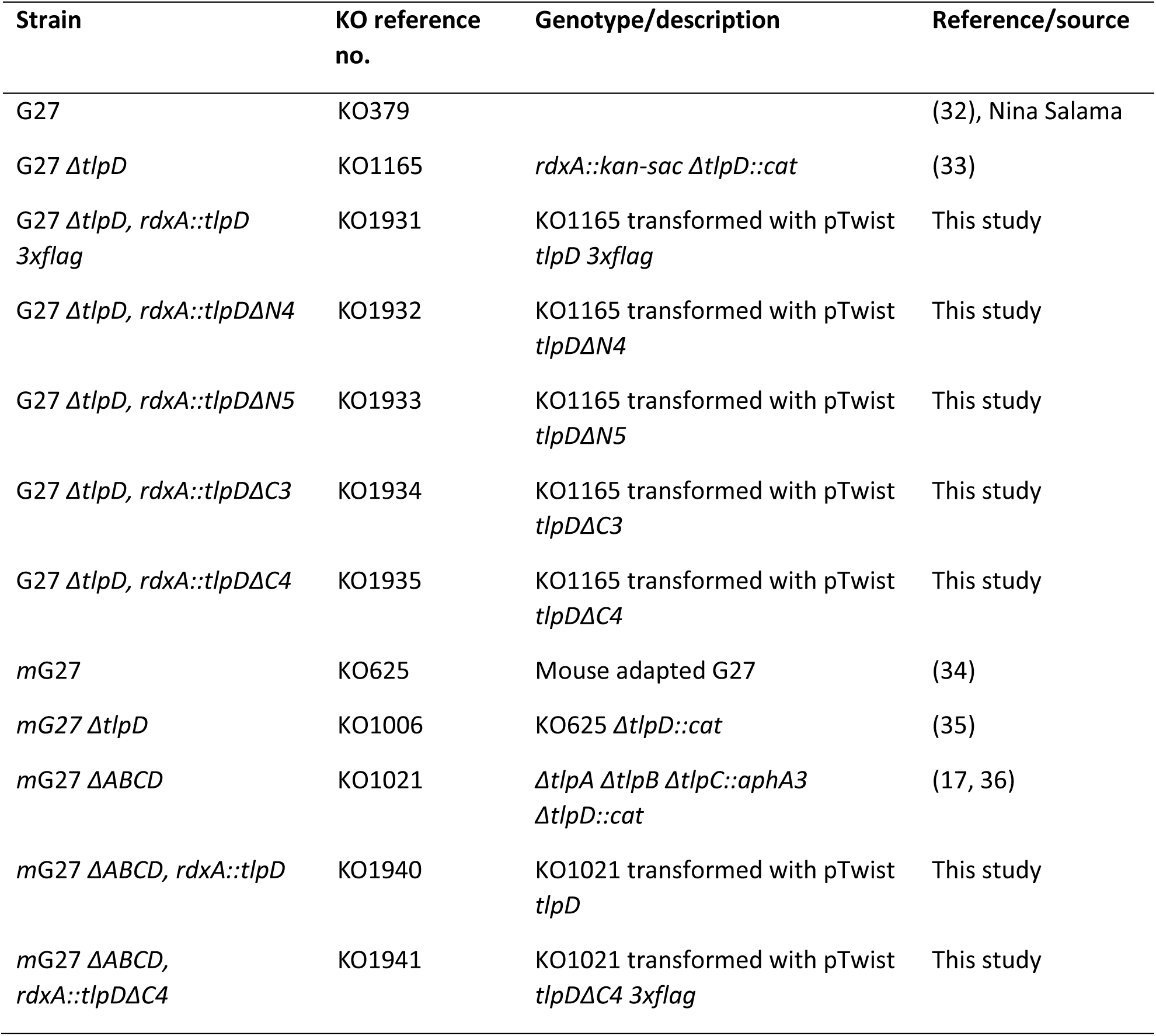
Bacterial strains used in this study.

| Strain | KO reference no. | Genotype/description | Reference/source |
| --- | --- | --- | --- |
| G27 | KO379 |  | (32), Nina Salama |
| G27 $\Delta tlpD$ | KO1165 | *rdxA::kan-sac \Delta tlpD::cat* | (33) |
| G27 $\Delta tlpD$, *rdxA::tlpD 3xflag* | KO1931 | KO1165 transformed with pTwist *tlpD 3xflag* | This study |
| G27 $\Delta tlpD$, *rdxA::tlpDΔN4* | KO1932 | KO1165 transformed with pTwist *tlpDΔN4* | This study |
| G27 $\Delta tlpD$, *rdxA::tlpDΔN5* | KO1933 | KO1165 transformed with pTwist *tlpDΔN5* | This study |
| G27 $\Delta tlpD$, *rdxA::tlpDΔC3* | KO1934 | KO1165 transformed with pTwist *tlpDΔC3* | This study |
| G27 $\Delta tlpD$, *rdxA::tlpDΔC4* | KO1935 | KO1165 transformed with pTwist *tlpDΔC4* | This study |
| mG27 | KO625 | Mouse adapted G27 | (34) |
| mG27 $\Delta tlpD$ | KO1006 | KO625 $\Delta tlpD::cat$ | (35) |
| mG27 $\Delta ABCD$ | KO1021 | $\Delta tlpA \Delta tlpB \Delta tlpC::aphA3 \Delta tlpD::cat$ | (17, 36) |
| mG27 $\Delta ABCD$, *rdxA::tlpD* | KO1940 | KO1021 transformed with pTwist *tlpD* | This study |
| mG27 $\Delta ABCD$, *rdxA::tlpDΔC4* | KO1941 | KO1021 transformed with pTwist *tlpDΔC4 3xflag* | This study |

### C-terminus truncated TlpDΔC4 retains interaction with CheV1 and CheW

TlpDΔC4 was next analyzed for its ability to fold. TlpD interacts with chemotaxis proteins including coupling proteins CheV and CheW (17). We therefore analyzed whether these interactions were retained for TlpDΔC4 using an approach previously used with *H. pylori* CheV1 and CheW, the bacterial two-hybrid (BACTH) analysis (16). Truncated *tlpD* was fused to the T18C fragment at the C-termini (T18C-*tlpD*, T18C-*tlpDΔC4*), as done for full-length *tlpD*, and then *tlpD*ΔC4 plasmids were analyzed with NT25 fragment plasmids with *cheV1* (*cheV1*-NT25) or *cheW* (*cheW*-NT25) (Fig. 3A) used previously (16). TlpDΔC4 was able to interact with both CheV1 and CheW (Fig. 3B) to levels that were significantly above the negative control. These levels are low but similar to ones obtained with full-length TlpD here (Fig. 3) and as reported previously (16). This result supports that the truncated TlpDΔC4 retains binding function with both CheV1 and CheW, suggesting the TlpDΔC4 MA domain is correctly folded.

**Figure 3.**
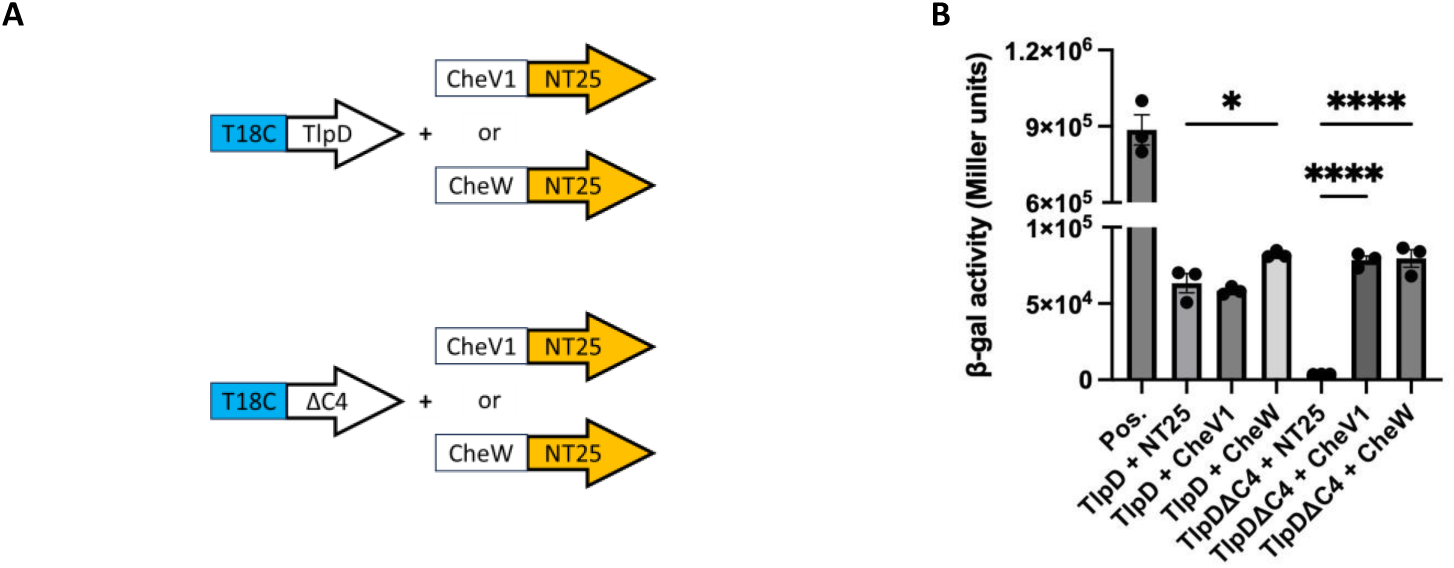
TlpD’s interaction with chemotaxis proteins is retained when the C-terminus is truncated. BACTH β-galactosidase (β-gal) assays of full-length and truncated (TlpDΔC4) TlpD with CheV1 and CheW. (A) Diagrams of constructs used. (B) β-gal assays. Positive (+) controls are T25-zip and T18-zip. Negative (−) controls are the respective TlpD construct with blank T25 plasmids. β-gal activity is calculated from three technical replicates with error bars indicating the standard error of the mean. *p < .05, ****p < .0001.

### TlpDΔC4 is unable to localize to the *H. pylori* cell pole

TlpD normally localizes to the *H. pylori* pole even when it is the only chemoreceptor present (17). We next analyzed the localization of TlpDΔC4 using immunofluorescence when it was expressed in either the presence of other chemoreceptors (TlpA, TlpB) or as the sole chemoreceptor. Western blot analysis of the receptorless *H. pylori* strain (Δ*ABCD*) complemented with TlpD shows protein expression (Fig. S4). Full-length TlpD localized to the pole in the presence or absence of the other chemoreceptors, as previously shown for a strain of *H. pylori* with natively expressed TlpD (17) (Fig. 4A and B). In contrast, TlpDΔC4 was diffuse throughout the bacterium’s cell cytoplasm (Fig. 4A). This cytoplasmic localization did not change when the other chemoreceptors were present (Fig. 4B). Altogether, using two different *H. pylori* strain backgrounds, these findings support the finding that the C-terminal portion of TlpD is important for polar localization.

**Figure 4.**
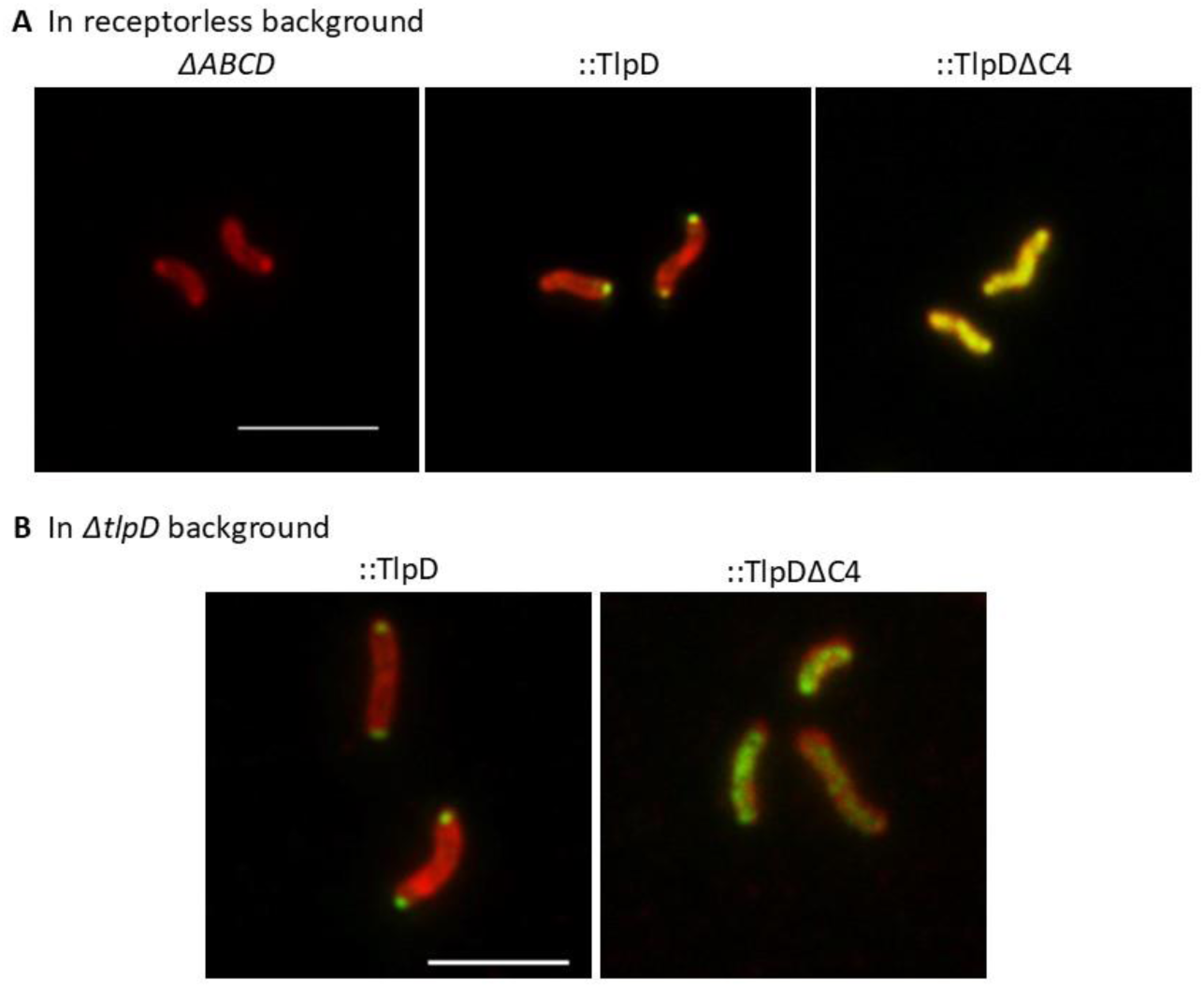
TlpD is polar when the C-terminus is intact and diffused when the C-terminus is disrupted. Immunolocalization of *H. pylori* whole cells producing full-length TlpD or truncated TlpDΔC4 protein. *H. pylori* was detected with chicken anti-*H. pylori* followed by goat anti-chicken conjugated to AlexaFluor 594 (red). TlpD proteins indicated above each image were detected with preabsorbed rabbit anti-TlpA22 (Δ*ABCD*, TlpD) or rat anti-FLAG (TlpD, ΔC4) followed by goat anti-rabbit or goat anti-rat conjugated to AlexaFluor 488 (green). Scale bar = 4μM. Images are representative of 3 biological replicates, with 20-50 bacterial cells analyzed for each sample.

### TlpDΔC4 alters localization of chemotaxis coupling proteins CheV1, CheV2, and CheV3 but not CheW

The above results show that TlpDΔC4 retains ability to interact with chemotaxis coupling proteins (Fig. 3) but mislocalizes (Fig. 4). The CheV and CheW coupling proteins normally are localized at the cell pole in *H. pylori* by interactions with the chemoreceptor complex (16, 17). We next analyzed how TlpDΔC4 would affect localization of these *H. pylori* chemotaxis signaling proteins, using strains that retained the TlpA and TlpB chemoreceptors and antibodies previously shown to be specific to each coupling protein (17, 36). Full-length TlpD expression resulted in all CheV and CheW proteins being entirely polar, with no detectable cytoplasmic staining, as previously shown with wild type *H. pylori* (17) (Fig. 5). In contrast, when TlpDΔC4 was expressed, CheV1, CheV2, and CheV3 displayed both polar and nonpolar staining, with CheV3 showing the most diffused cytoplasmic signal (Fig. 5). CheW, in comparison, showed only polar staining. These results further support that TlpDΔC4 retains the ability to interact with chemotaxis coupling proteins and can retain them in the cytoplasm.

**Figure 5.**
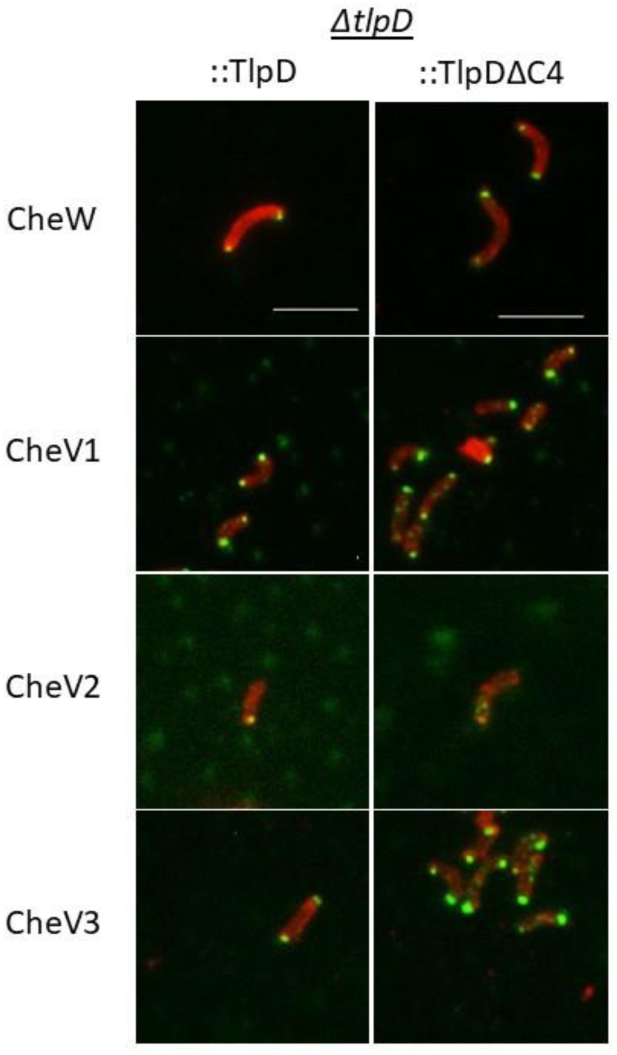
Localization of CheW and CheV coupling proteins in *H. pylori* in the presence of TlpDΔC4. Immunolocalization of *H. pylori* chemotaxis proteins CheW, CheV1, CheV2, and CheV3 in whole cells producing full-length (TlpD-FLAG) or truncated (TlpDΔC4-FLAG) protein. *H. pylori* was detected with chicken anti-*H. pylori* followed by goat anti-chicken conjugated to AlexaFluor 594 (red). Individual chemotaxis proteins indicated on the left were detected with preabsorbed guinea pig anti-CheW, rabbit anti-CheV1, rabbit anti-CheV2 or rabbit anti-CheV3 followed by goat anti-guinea pig or goat anti-rabbit conjugated to AlexaFluor 488 (green). Scale bar = 4μM. Images are representative of 3 biological replicates, with 20-50 bacterial cells analyzed for each sample.

### TlpDΔC4 does not confer soft agar migration in *H. pylori*

To explore TlpDΔC4 function, chemotaxis ability of strains expressing TlpDΔC4 was analyzed using a field-standard chemotaxis assay, migration in soft agar. In this assay, bacteria are inoculated at a single point, and if motile and chemotactic, migrate outwards to form expanded colonies. Using strains that express the *rdxA* copy of *tlpD* as their only chemoreceptor, we found that full-length TlpD was able to confer spreading to diameters that were significantly elevated compared to the parent strain lacking all chemoreceptors (*ΔABCD*) (Fig. 6A and C). These findings suggest that complementation of wild-type *tlpD* in the *rdxA* locus is partially sufficient to restore spreading motility in *H. pylori*. The *tlpD*ΔC4 complemented strain, in contrast did not spread even after seven days (Fig. 6B and D). This finding suggests that truncation of the C-terminus of TlpD results in a chemoreceptor that is insufficient to restore spreading. These findings suggest that TlpDΔC4 may be unable to establish a working chemotaxis system.

**Figure 6.**
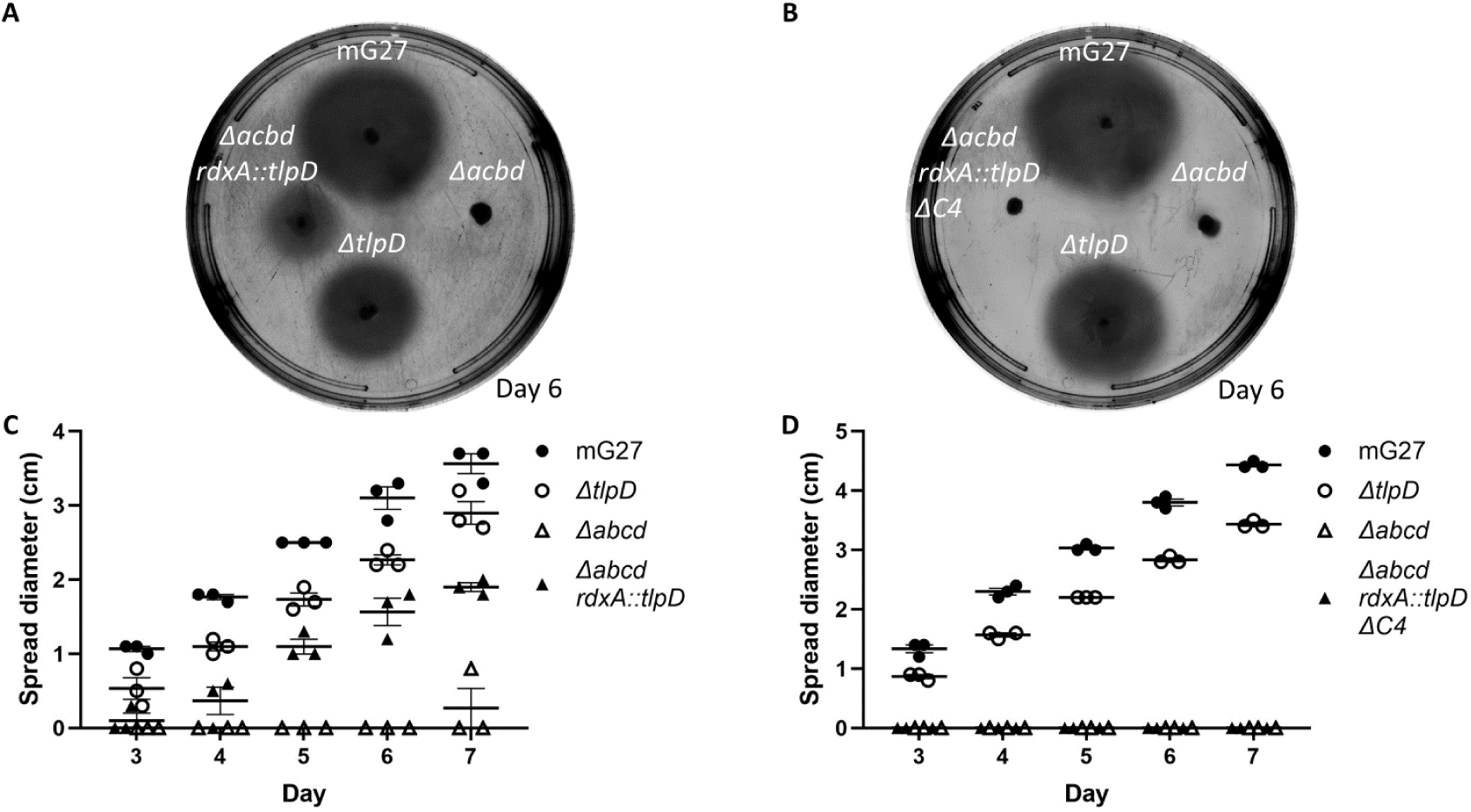
*H. pylori* that express TlpDΔC4 as the sole chemoreceptor do not spread on soft agar. (A) *H. pylori* wild-type mG27 and isogenic mutants were analyzed for chemotaxis by soft agar spreading. Strains were stabbed onto soft agar and spreading was measured after 3 days. Strains included are the parent *H. pylori* lacking all chemoreceptors (*ΔABCD,* called *Δabcd* here), and isogenic versions complemented with a full-length TlpD (*rdxA::tlpD*) (A) or with TlpDΔC4 (*rdxA::tlpD ΔC4*) (B). Plates are representative of 3 biological replicates, with three technical triplicates. Error bars represent the standard error of the mean.

## Discussion

Our experiments with a truncated version of the cytoplasmic protein TlpD revealed that TlpD polar localization requires its C-terminal 45 amino acids. We reached this conclusion using immunofluorescence of FLAG-tagged full length TlpD or FLAG-tagged TlpDΔC4. Full length TlpD-FLAG localized to the pole as reported for TlpD lacking the FLAG (17), suggesting the FLAG tag did not disrupt any localization signals, as seen with a previous V5 tag (37). TlpDΔC4-FLAG, in contrast, did not display polar localization, suggesting that the C-terminal 45 amino acids are critical for localization directly or for allowing TlpD to adopt a form that can localize. Our analysis furthermore suggests that the TlpDΔC4 protein can fold, based on its ability to maintain interactions with CheV proteins in *H. pylori* and CheV1 and CheW in the BACTH assays. Interestingly, CheV proteins mislocalized to the cytoplasm more than CheW did when TlpDΔC4 was present, suggesting that TlpD might have an interaction preference for CheV proteins versus CheW, as suggested by previous BACTH studies of TlpD (16) and seen in *Campylobacter jejuni* chemoreceptors (38). Overall, these findings support the new idea that TlpD requires its C-terminal 45 amino acids to properly localize.

Our search to identify features within the C-terminal 45 amino acids of TlpD revealed no apparent hits. The AlphaFold structure described here suggests that this region adopts an extended 43 amino acid alpha helix that sits alongside the MA and CZB domains on one side, leaving the other side exposed on the surface (Fig. 1A). Similarly, it is possible that the N and C-termini of TlpD interact with one another, based on the idea that it was difficult to truncate either region. Thus, this region is clearly important but not yet clear whether it plays a structural, regulatory or protein-protein interaction function.

A potential function of this C-terminal stretch is in promoting protein-protein interactions. Previous work showed that TlpD’s polar localization was dynamic, with TlpD residing at the pole under conditions that correlated with high energy, and diffuse in conditions correlated with low ATP (25, 37). Besides the chemotaxis signaling proteins, several other proteins have been identified to interact with TlpD. Yeast two-hybrid analyses identified TlpD interactors as acetone carboxylase (AcxC, HP0599) (39) and sialic acid binding protein (HP0721) (40, 41). Two additional proteins, aconitase (AcnB) and catalase (KatA) were shown to interact with TlpD based on pull downs from *H. pylori* extracts (37). AcnB, KatA, and AcxC were further studied for their ability to localize TlpD to the pole by analyzing TlpD position in mutants lacking the genes for each protein. Loss of *katA* resulted in TlpD being less polar, and observed throughout the cell body (37). Whether TlpD interacts directly with KatA and via which regions is not known. This work however corroborates that TlpD is polarly localized in a manner that is regulated and via specific protein-protein interactions beyond the chemotaxis signaling network.

TlpD plays a critical role in building the chemotaxis protein arrays. Previous work had shown that TlpD as the sole chemoreceptor localizes to the pole, and is sufficient to localize the main components of the chemotaxis signaling complex—CheA, CheW, CheV1 and CheV3 to the cell pole too (17). Furthermore, work with *H. pylori* that expressed only one transmembrane chemoreceptor, either TlpA or TlpB, and lacked TlpD, mislocalized all three CheV proteins (42). These studies suggest TlpD plays a critical role in promoting chemotaxis proteins to localize to the pole. Our localization studies showed that even when using an *H. pylori* strain that retained TlpA and TlpB, two transmembrane chemoreceptors, the TlpDΔC4 was not at the pole and mislocalized CheV proteins (Fig. 5). These findings suggest that transmembrane chemoreceptors of *H. pylori* are not sufficient to maintain a distinctly polar chemotaxis complex, but rather require polar TlpD.

These findings provide insight into the workings of cytoplasmic chemoreceptor proteins. The cytoplasmic chemoreceptors that localize to the cell pole do so by colocalization with transmembrane chemoreceptors (43–45), and in some cases switch between a diffused cytoplasmic protein and a distinct polar topology (37, 46). Cytoplasmic chemoreceptors presumably utilize similar mechanisms as those used by other proteins to drive polar localization including stochastic self-assembly, interaction with polar anchor proteins, or interactions with plasmid partitioning homologues (47). TlpD contains a CZB domain, and our findings may be specific to these types of chemoreceptors. One candidate for comparative analysis is the CZB-containing cytoplasmic chemoreceptor McpA (NCBI Ref Seq: WP_000475246.1) from the enteric pathogen *Salmonella enterica*. McpA is a 352 amino acid chemoreceptor protein with an identified MA domain, a C-terminal CZB domain, and an N-terminal coiled coil (48, 49). *S. enterica* that lacks seven of its nine chemoreceptors, retaining only McpA and another of unknown function called Tip, fail to spread on soft agar (50, 51). McpA was not the focus of these previous studies, however the findings do suggest this protein is not sufficient to confer a spreading phenotype in *S. enterica*. The alignment of full length TlpD and McpA shows 33% amino acid identity over the entire protein, and the C-terminal domains share 29% amino acid identity. Moreover, McpA retains the conserved CZB motif (**H**xx[WFYL]x_21-28_**C**x[LFMVI]Gx[WFLVI]x_18-27_**H**xxx**H**) at its C-terminus (amino acids 246-303) (23). The localization of McpA is not yet known however and will be interesting for future studies.

In sum, our work shows that TlpD supports building of a functional chemosensory complex at the pole and relies on the sequences at its C-terminal end to accomplish this function. One model, consistent with our data, is that the C-terminus of TlpD interacts with an as-yet-identified polar landmark protein, to bring TlpD to the pole, possibly via catalase. Once there, TlpD plays a critical role in nucleating chemosensory array formation. Testing this model and exploring whether other cytoplasmic chemoreceptors have similar localization sequences is an exciting area for future work.

## Materials and Methods

### Bacterial strains and growth conditions

All *H. pylori* and *E. coli* strains used in this study are listed in Tables 1 and 2. *H. pylori* was grown on Columbia Horse Blood Agar (CHBA) containing 5% (vol/vol) defibrinated horse blood (Hemostat Labs, Davis, CA), 0.2% (wt/vol) β-cyclodextrin (TCI), 5 μg/mL cefsulodin (Sigma Aldrich), 50 μg/mL cycloheximide (GoldBio), 2.5 U/mL polymyxin B (Sigma Aldrich) and 10 μg/mL vancomycin (GoldBio). For the selection of *H. pylori* mutants, 13 μg/mL chloramphenicol, 15 μg/mL kanamycin (Fisher BioReagents), or 25 μg/mL erythromycin (GoldBio) was used. For liquid cultures, *H. pylori* was grown with brucella broth containing 10% heat-inactivated fetal bovine serum (Gibco) (BB10). Cultures were incubated at 37°C in microaerobic conditions (5% O_2_, 10% CO_2_, and 85% N_2_). *E. coli* was grown on LB media containing 100 μg/mL ampicillin or 30 μg/mL kanamycin (Fisher BioReagents) and incubated at 37°C unless otherwise stated.

### Strategy for creation of truncated TlpD

To construct truncated versions of G27 TlpD (Uniprot:B5Z6X0), Protein CCD (29) was used to identify predicted secondary structures, features, and domains within the protein and served as the basis for selecting truncation sites that avoid secondary structures and predicted or known regions. TlpD N-terminal truncations at amino acids 53 (ΔN4) and 104 (ΔN5), were selected to sit in between secondary structures. The TlpD C-terminus truncations at amino acids 322 (ΔC3) and 388 (ΔC4) were selected to abolish or retain the CZB domain without interrupting the canonical MA domain, respectively. Additionally, 200bp upstream of *tlpD* was included to capture any transcriptional regulatory sites and sequences encoding a 3xFLAG tag (DYKDDDDK-DYKDDDDK-DYKDDDDK) was added to the C-terminus of TlpD. Lastly, an erythromycin resistance gene and its promoter were added for positive selection, downstream of *tlpD*. This entire sequence was flanked by 500bp upstream and downstream of the non-native locus, *rdxA*, for chromosomal integration in *H. pylori*. pTwist Amp High Copy plasmids with ampicillin resistance were ordered from Twist Bioscience that contained these *tlpD* cassette sequences (File S2). These plasmids were then transformed into electrocompetent *E. coli* DH10B cells using ampicillin selection followed by single colony purification.

### Creation of G27 and mG27 *tlpD* complement mutants

For transformation into *H. pylori*, plasmids containing *tlpD* sequences were purified from *E. coli* DH10B cells using the QIAprep Spin Miniprep Kit (Qiagen) and added to a suspension of BB10 and 1-day old *H. pylori* G27 *ΔtlpD* or *ΔtlpA ΔtlpB ΔtlpC ΔtlpD* cells from a CHBA plate (52). Transformants were selected by growth on CHBA containing 25 μg/mL erythromycin followed by single colony purification. All mutations were verified using PCR with primers that flank *rdxA* (*rdxA* Fwd and Rev) (Table S1).

### Western blot

The expression of modified TlpD was observed by western blot. Liquid overnights of *E. coli* were grown and used to prepare lysate samples. 1-to-2-day old *H. pylori* grown on CHBA solid media containing erythromycin were resuspended in 1X Phosphate-Buffered Saline (PBS) and used to prepare lysate samples. To prepare the lysate, samples were back diluted to OD_600_ = 0.7 in 200 μL. 6X SDS PAGE sample buffer was added to the mixture at 1X final concentration and 1X PBS was added to 200 μL. Next, samples were heated at 95-100°C for 10 minutes then placed on ice. Samples were electrophoresed on 10% or 12% SDS-PAGE gels then semi-dry transferred to polyvinylidene (PVDF) membranes using a trans-blot SD semi-dry transfer cell (Bio-Rad). Relative protein loading was visualized by staining the membrane with Direct Blue 71 (DB71) stain containing 0.008% DB71 (Sigma Aldrich) in 40% EtOH and 10% acetic acid prior to antibody binding. PVDF membranes were incubated with a 1:20,000 dilution of rat anti-FLAG monoclonal antibody (Novus Biological, NBP1-06712) or a 1:4,000 to 1:10,000 dilution of rabbit anti-TlpA22 antibody (33) diluted in 5% BSA at 4°C overnight. Membranes were next incubated with a 1:20,000 dilution of horseradish peroxidase-conjugated goat anti-rat (R&D Systems) or a 1:10,000 dilution of goat anti-rabbit (Santa Cruz Biotechnology) secondary antibody. For visualization, SuperSignal West Pico PLUS Chemiluminescent Substrate (Thermo Scientific) was added at a 1:1 ratio for 30 to 120 seconds with shaking at room temperature. Blots were then imaged using a Bio-rad Chemi-doc.

### Soft-agar motility assay

*H. pylori* spreading motility was carried out as previously described (17, 53–55). Briefly, soft agar plates were made with brucella broth and 0.35% bacto agar (BD). Soft agar was cooled to 50°C before 2.5% heat-inactivated fetal bovine serum, 8 μg/mL amphotericin B, 5 μg/mL cefsulodin, 50 μg/mL cycloheximide, 2.5 U/mL polymyxin B, 5 μg/mL trimethoprim (Sigma Aldrich), and 10 μg/mL vancomycin were added. Plates set for 3 days at room temperature before use. 1–2-day old *H. pylori* strains from CHBA plates were inoculated halfway into the agar using a pipet tip. Plates were incubated right side up in the 37°C microaerobic chamber and migration diameter was measured daily. mG27, mG27 *ΔtlpD*, and mG27 *ΔtlpA ΔtlpB ΔtlpC ΔtlpD* were used as controls. Statistical analysis was done on GraphPad Prism v9.4.1.

### Immunofluorescence

Immunolocalization of *H. pylori* chemotaxis proteins TlpD, CheW, CheV1, CheV2, and CheV3 was performed as previously described (17, 36). Briefly, 1-day old *H. pylori* grown on Columbia Horse Blood Agar (CHBA) under microaerobic conditions was harvested and resuspended in 1mL of 1X PBS. 20 μL of the cell suspension was placed on a Poly-Lysine glass slide (Fisher Scientific) and fixed with 0.075M NaPO_4_ adjusted to pH7.4, 0.0025M NaCl and 2% paraformaldehyde (EMS) solution then permeabilized in 3% Bovine Serum Albumin (BSA) (Millipore), 1% Saponin (Calbiochem), 0.01% Triton X-100 and 0.02% Na Azide (Sigma Aldrich) resuspended in 1X PBS. The samples were blocked with 3% BSA and 0.01% Triton X-100 resuspended in 1X PBS solution. Samples were next treated with a 1:500 dilution of chicken anti-*H.pylori* (Agrisera AB), a 1:2,000 dilution of rat anti-FLAG (Novus Biological, NBP1-06712), a 1:200 of preabsorbed or non-preabsorbed rabbit anti-TlpA22, or a 1:50 dilution of preabsorbed rabbit anti-CheV1 (36), anti-CheV2, anti-CheV3 (J. Castellon, P. Lertsethtakarn, and K.M. Ottemann, unpublished), or guinea pig anti-CheW (17) diluted in permeabilization solution (described above). The anti-TlpA22, anti-CheV1, anti-CheV2, and anti-CheV3 antibodies were previously shown to be specific to each coupling protein (17, 36). The samples were then treated with a 1:500 dilution of goat anti-chicken Alexa Fluor 594 (Invitrogen, A11042) and goat anti-rat Alexa Fluor 488 (ThermoFisher, A48262) or a 1:300 dilution of goat anti-rabbit Alexa Fluor 488 (Invitrogen, A11008) or goat anti-guinea pig Alexa Fluor 488 (Invitrogen, A11073) diluted in permeabilization solution. The samples were washed with blocking solution (described above) 3 times, a drop of Vectashield (Vector Laboratories) was added to the cells and then sealed with coverslips. Fluorescence microscopy, using the Nikon Eclipse E600 was used to visualize immunofluorescent cells at 100X objective with oil immersion. Texas Red and Fitc/GFP filters were used to separately view and capture emission spectra from Alexa Fluor 594 (red) and 488 (green), respectively. Images were captured using a Hamamatsu ORCA-285 digital camera equipped to the microscope. Fluorescent images were pseudo colored using ImageJ and merged using Adobe Photoshop version 24.1.0. Scale = 4 μm. Images are representative of 3 biological replicates.

### Creation of Bacterial Two-hybrid (BACTH) plasmids and BACTH analysis

Synthetic plasmid DNA from *E. coli* DH10B was extracted using the Qiaprep Mini Kit (Qiagen). *H. pylori* G27 full-length *tlpD* and truncated *tlpDΔC4* constructs were amplified using PCR with primers that amplify each sequence (*tlpD* pUT18 Fwd and Rev, *ΔC4* pUT18 Fwd and Rev) (Table S1). The PCR products encoded PstI and SacI restriction sites which were digested with Pst1-HF (NEB) and SacI-HF (NEB) restriction enzymes. BACTH plasmids pUT18 and pUT18C (J. Gober, UCLA) (31) were similarly digested followed by phosphatase treatment using rSAP following the manufacturer’s suggested protocol (NEB). Treated PCR products and plasmids were purified using the GFX PCR DNA and Gel Band Purification Kit (Cytiva). The purified products were ligated using T4 DNA ligase following the manufacturer’s suggested protocol (NEB) and subcloned into electrocompetent *E. coli* XL1-Blue cells and ampicillin resistant colonies were selected. All constructs were confirmed using PCR with the same primers used to amplify *tlpD* or *tlpDΔC4* (Table S1) and DNA sequencing (Azenta).

Recombinant plasmids encoding the *cheV1*-NT25 (16), *cheW*-T25 (16), T18C-*tlpD*, T18C-*tlpDΔC4* and were co-transformed, at 40 ng each, into electrocompetent BTH101 and plated on LB containing 40 μg/mL X-gal (5-bromo-4-chloro-3-ondolyl-beta-galacto-pyranoside) (GoldBio), 0.5 mM IPTG (Isopropyl β-d-1-thiogalactopyranoside) (Fisher BioReagents), 100 μg/mL ampicillin, and 50 μg/mL kanamycin. As positive controls, T25-zip and T18-zip plasmids were co-transformed into BTH101. As negative controls, each *tlpD* recombinant plasmid was co-transformed with a pKNT25 blank BACTH plasmids. For interaction assays, colonies including controls grown on LB containing X-gal, IPTG, ampicillin, and kanamycin at the above-mentioned concentrations were used to make overnights in LB with ampicillin and kanamycin at the above concentrations. The following day, 20 μL of each overnight was dropped onto LB plates containing X-gal, IPTG, ampicillin, and kanamycin as above. These spots were allowed to dry at room temperature before the plates were incubated at 30°C for 24 hours. Images were captured using an iPhone 14 Pro and a white light transilluminator (Fisher Scientific). Images are representative of 3 biological replicates.

### β-Galactosidase Assay

To quantify BACTH interactions, β-galactosidase activity was measured as previously described (16, 31). Briefly, transformants were grown in LB broth with ampicillin, kanamycin, and IPTG at the above concentrations at 30°C overnight. Overnight cultures were back diluted to OD_600_= 0.1 in LB broth with ampicillin, kanamycin, and IPTG then grown at 30°C until OD_600_ 0.3-0.7 was reached. 100 μL of each culture was mixed with 900 μL cold Z-Buffer (0.06M Na_2_HPO_4_.7H_2_O, 0.04M NaH_2_PO_4_.H_2_O, 0.01M KCl, 0.001M MgSO_4_.7H_2_O, adjusted to pH 7.0 and 0.05M β-mercaptoethanol was added fresh before use) in triplicate and mixed by inversion. Next, 50 μL of 0.1% SDS and 100 μL of chloroform was added to each sample followed by a 10 second vortex. The samples and fresh 4mg/mL ortho-Nitrophenyl-β-galactoside (ONPG) (Thermo Scientific) (in 0.1M Phosphate Buffer solution (0.06M Na_2_HPO_4_.7H_2_O and 0.04M NaH_2_PO_4_.H_2_O adjusted to pH 7.0) were incubated at 30°C for 5 minutes. 200 μL ONPG solution was added to samples, vortexed, and time was recorded. The samples were incubated at 30°C until a yellow color appeared. The reactions were stopped by adding 500 μL Stop solution (1M Na_2_CO_3_), time recorded, and samples centrifuged at room temperature and max speed for 5 minutes before OD_420_ was measured. Miller units were calculated to determine β-galactosidase activity using the following formula: 1000 x (OD_420_ / [time * vol. culture * OD_600_]) (56). β-gal activity is one biological replicate with three technical replicates per sample. Statistical analysis was done on GraphPad Prism v9.4.1.

## Acknowledgements

We thank Fitnat Yildiz, Carrie Partch, and Chad Saltikov for discussion and support. We also thank UCSC STEM Diversity for their support, particularly Yulianna Ortega and Xingci Situ. We are grateful to past and current Ottemann lab members for their continued support throughout this project, particularly Arturo Valdez and Christina Yang. The described project was supported by the National Institute of Allergy and Infectious Diseases (NIAID) grant RO1AI116946 to K.M.O. The funders had no role in study design, data collection, and interpretation, or decision to submit the work for publication.

## Supporting information

**Figure S1.**
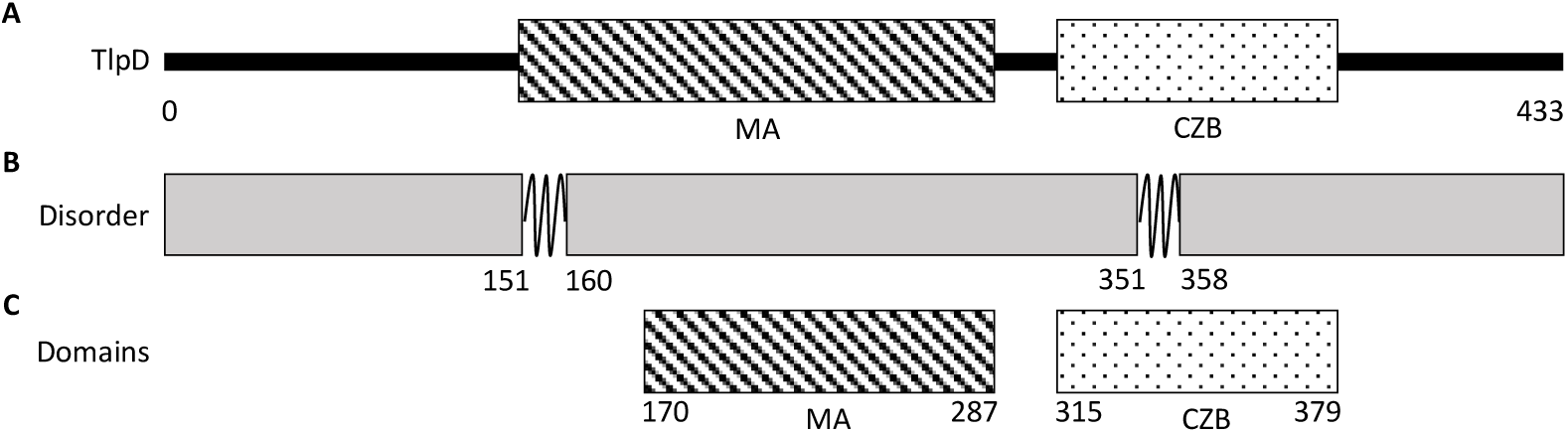
Summary of Predicted secondary structures and domains of TlpD. (A) Full-length TlpD for reference. (B) Predicted TlpD globular regions in grey and the unstructured regions shown as twists. (C) Predicted TlpD domains with the two highest scores, the MA and CZB domain.

**Figure S2.**
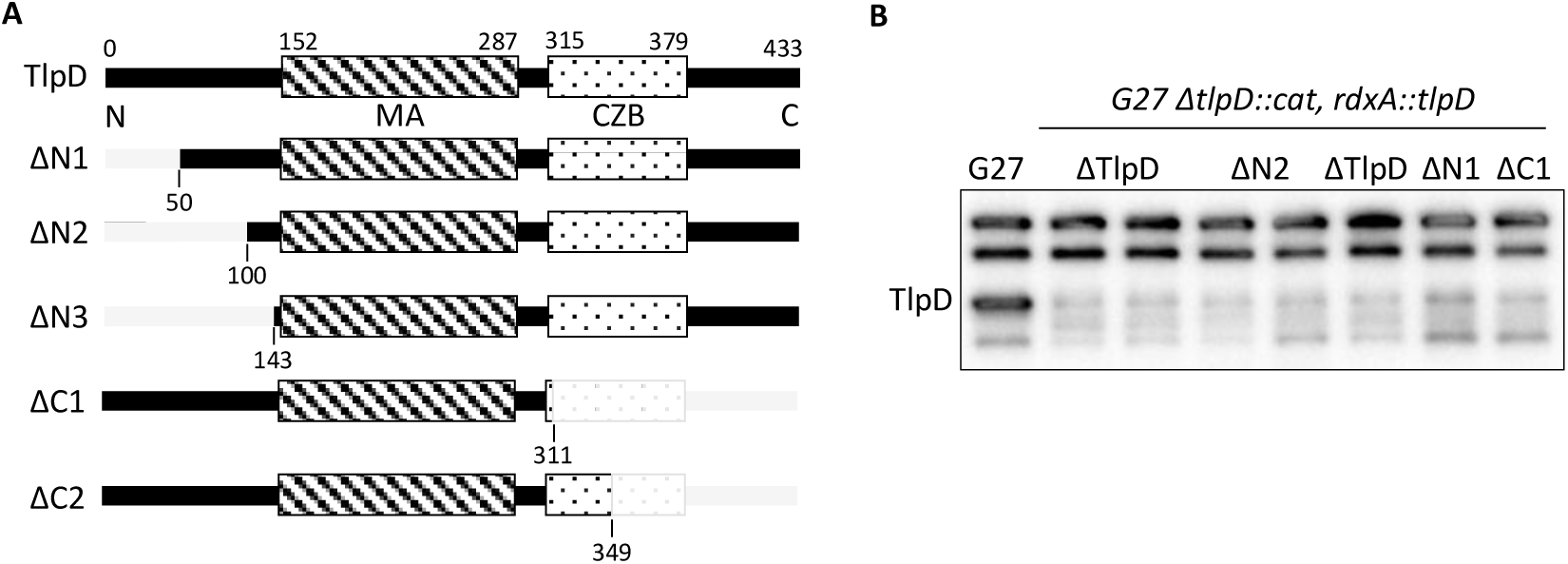
First generation of TlpD truncation constructs and expression in *H. pylori*. (A) Modified versions of *H. pylori* TlpD with domains and truncation sites (grey). TlpD is the full-length version of the protein at 433 amino acids, ΔTlpD are negative background controls, ΔN1 is truncated at residue 50, ΔN2 is truncated at residue 100, ΔN3 is truncated at residue 143, ΔC1 is truncated at residue 311, and ΔC2 is truncated at residue 349. (B) Western blot of *H. pylori* G27 strains with complemented *tlpD* constructs were analyzed from whole-cell lysates with an anti-TlpA-22 antibody (33) which recognizes the conserved MA domain of all chemoreceptors. The expected size of wild-type TlpD is 48 kDa, ΔN1 is 42.9 kDa, ΔN2 is 37.4 kDa, and ΔC1 is 34 kDa. Western blots are representative of 3 biological replicates.

**Figure S3.**
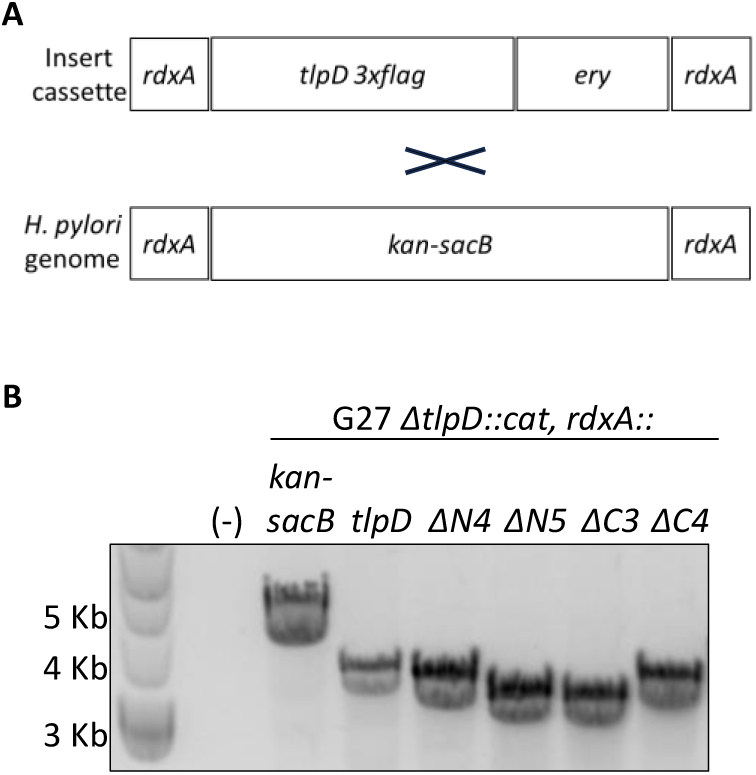
*tlpD* sequence layout for transformation into the *H. pylori* genome. (A) Design of *tlpD* insert cassette for transformation into *H. pylori*. This cassette contains the *tlpD* sequence, a *3xflag* tag and *erythromycin* (*ery*) resistance sequences that are flanked by *rdxA* homologous regions for integration into the *H. pylori* genome at the *rdxA* locus. (B) PCR of erythromycin resistant *H. pylori* isolates using primers that flank the *rdxA* locus. The nomenclature above each band indicates the amplified product. (−); PCR no template control. *kan-sacB*; kanamycin resistance and sucrose sensitivity selection marker present in the *H. pylori* transformation background and serves as a negative control. *tlpD*, ΔN4, ΔN5, ΔC3, ΔC4 are the versions of *tlpD* on the cassette that were PCR amplified.

**Figure S4.**
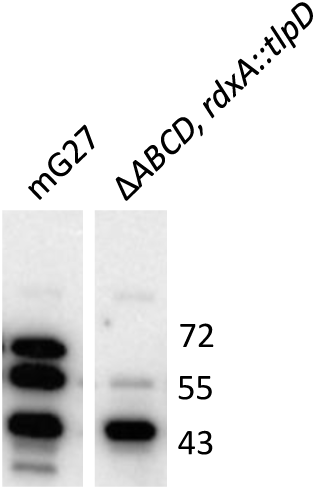
TlpD is produced when complemented into *H. pylori* that lacks all chemoreceptors. Western blot of *H. pylori* mG27 *Δabcd* expressing complemented WT *tlpD* was analyzed from whole-cell lysates with an anti-TlpA22 antibody which recognizes the MA domain. The strains analyzed are indicated at the top. Marker sizes are given in kilodaltons on the right. The expected size of TlpD is 48 kDa. Western blot is representative of 2 biological replicates.

**Table S1.**
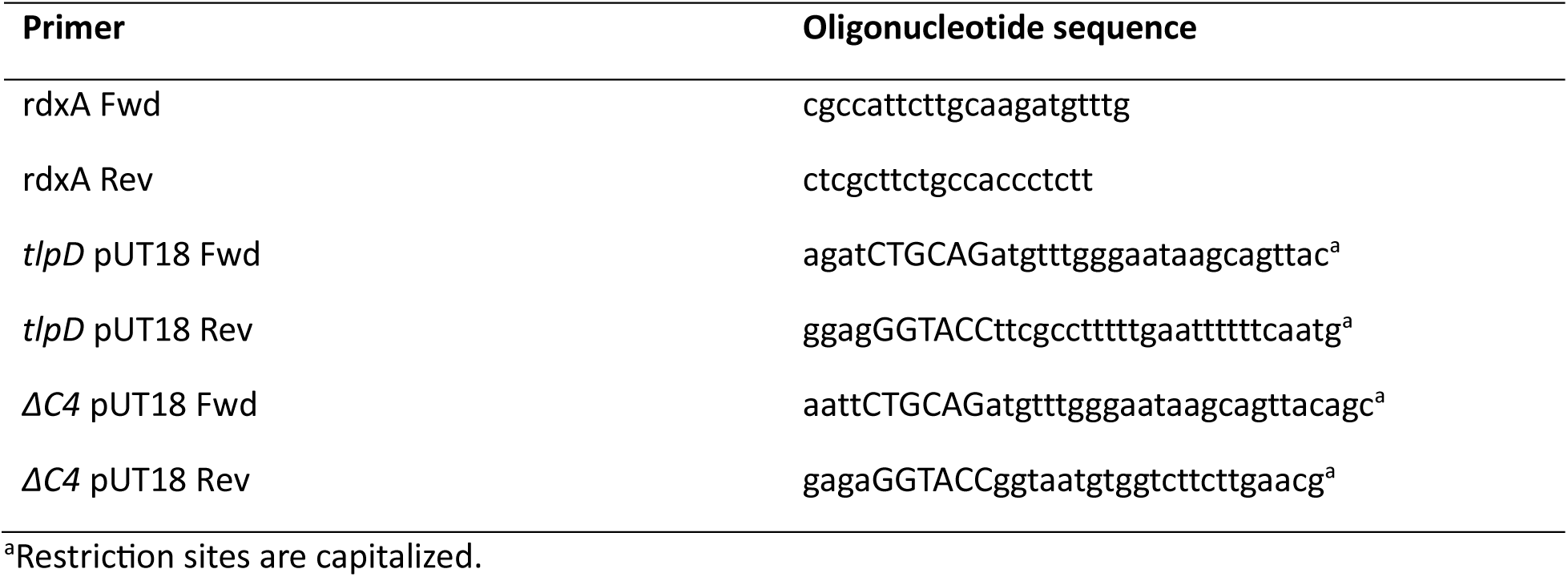
Primers used in this study.

File S1. TlpD secondary structure bioinformatic analysis.

File S2. *tlpD* plasmid sequences.

